# Production of interspecies somatic/pluripotent heterokaryons using polyethylene glycol (PEG) and selection by imaging flow cytometry for the study of nuclear reprogramming

**DOI:** 10.1101/655621

**Authors:** M. Cristina Villafranca, Melissa R. Makris, Maria Jesus Garrido Bauerle, Roderick V. Jensen, Willard H. Eyestone

## Abstract

Fusion of somatic cells to pluripotent cells such as mouse embryonic stem (ES) cells induces reprogramming of the somatic nucleus, and can be used to study the effect of *trans*-acting factors from the pluripotent cell on the somatic nucleus. Moreover, fusion of cells from different species permits the identification of the transcriptome of each cell, so the gene expression changes can be monitored. However, fusion only happens in a small proportion of the cells exposed to fusogenic conditions, hence the need for a protocol that produces high fusion rate with minimal cell damage, coupled with a method capable of identifying and selecting fusion events from the bulk of the cells. Polyethylene glycol (PEG) is a polymer of repeated ethylene oxide units known to induce cell fusion within a certain range of molecular weight. Here, we describe a method to induce formation of bi-species heterokaryons from adherent mammalian cells, which can then be specifically labeled and selected using live cell immunostaining and a combination of imaging and traditional flow cytometry. First, we tested several PEG-based fusion conditions to optimize a protocol to consistently produce both mouse NIH/3T3 fibroblast and primary bovine fetal fibroblast (bFF) homokaryons. Initially, we obtained 7.28% of NIH/3T3 homokaryons when using 50% PEG 1500. Addition of 10% of DMSO to the PEG solution increased the percentage of NIH/3T3 homokaryons to 11.71%. In bFFs, treatment with 50% PEG 1500 plus 10% DMSO produced 11.05% of homokaryons. We then produced interspecies heterokaryons by fusing mouse embryonic stem (mES) cells to bFFs. To identify bi-species fusion products, heterokaryons were labeled using indirect immunostaining in live cells and selected using imaging (Amnis ImageStream Mark II) and traditional (BD FACSAria I) flow cytometry. Heterokaryons selected with this method produced ES cell-like colonies when placed back in culture. The method described here can also be combined with downstream applications such as nucleic acid isolation for RT-PCR and RNA-seq, and used as a tool to study cellular processes in which the effect of *trans*-acting factors is relevant, such as in nuclear reprogramming.

## INTRODUCTION

Cell fusion is a process that involves combining the cytoplasmic membrane of two or more cells to form one single multinucleated cell[1]. Nuclear reprogramming to pluripotency can be achieved by fusion of somatic cells to pluripotent cells such as embryonic stem (ES) cells[2–4], embryonal germ (EG) cells[5], or embryonal carcinoma (EC) cells[6], suggesting the reprogramming activity of the pluripotent cell is predominant over the gene expression pattern of the somatic cell. The nuclei of somatic cells fused to ES cells reprogram faster (1 to 2 days) and with greater efficiency (up to 70%) than cells reprogrammed by transcription factor induction[1]. Fused cells can become hybrids or heterokaryons depending if their nuclei fuse or remain intact, respectively. Gene expression changes in heterokaryons happen in the absence of cell division, without genomic mixing or transgenes, which makes cell fusion a powerful tool to study early modifications in gene expression. Moreover, the use of interspecies heterokaryons allows the discrimination of the transcriptome of each fusion partner, thus facilitating the screening of factors provided by the pluripotent cell, and the changes in gene expression induced in the somatic cell. The cell fusion approach has already provided valuable information about reprogramming[7] and differentiation[8] mechanisms in human and mouse cells. However, both processes, which constitute of the foundations of regenerative medicine and developmental biology in mouse and human are, to date, still not fully understood or optimized[9]. Moreover, in other domestic and biomedically relevant species where pluripotent cell lines are not yet available[10,11], the interspecies heterokaryon approach could provide new insights as to which genes and mechanisms are essential in the establishment and maintenance of pluripotency in these species.

Despite technological advances in cell sorting and transcriptome analysis, cell fusion has yet to be used to its full potential, and protocols designed specifically for the study heterokaryons in a nuclear reprogramming context are not easily available. Due to its simplicity and low cost, polyethylene glycol (PEG) is usually the method of choice to induce fusion in a laboratory setting. PEG is a polyether of repetitive ethylene oxide units that can be classified according to the number of ethylene oxide units or their approximate molecular weight (e.g. PEG-32 or PEG 1500, respectively), and is used widely as a drug delivery agent and in cosmetics [12,13]. PEG in the molecular weight range of 1000 – 6000 has been described to induce cell fusion *in vitro*[14]. To date, most published work on PEG-induced mammalian cell fusion uses a combination of PEG in the molecular weight range of 1000 – 3700 [15–21]. Fusion can only be induced in a small fraction of the cells exposed to fusogenic conditions, and different cell types might require modifications of the method used in terms of PEG molecular weight, concentration, and/or time[14].

Here, we present a method to produce bi-species pluripotent/somatic heterokaryons, as well as same species homokaryons, from mammalian adherent cells growing on monolayers, which can be selected using indirect immunostaining in live cells and a combination of imaging and traditional flow cytometry. First, we produced a method to consistently produce a high number of viable multinucleated homokaryons from mouse fibroblasts (NIH/3T3 line) or primary bovine fetal fibroblasts by exposing cellular monolayers to PEG additioned with 10% DMSO; with this method, we obtained 11.71% and 11.05% of binucleated cells, respectively. Next, we used these conditions to produce interspecies heterokaryons formed by the fusion of co-cultured mouse embryonic stem (mES) cells (C57BL/6 line) to bovine fetal fibroblasts (bFFs), and identified them by indirect immunostaining in live cells, using fluorescent antibodies targeting mES cell-specific marker SSEA-1 and bFF-specific marker CD44, as well as nuclear staining using Hoechst. The population of double positive heterokaryons was identified first using the ImageStream Mark II imaging flow cytometer, and then selected using FACSAria I. Imaging flow cytometry was an essential step to identify the small heterokaryon cell population (∼0.1%). Because of its specificity, the heterokaryons selected with this method were suitable for downstream applications such as nucleic acid isolation for PCR and RNA-seq, or long-term culture to produce clonal hybrid colonies. Our selection protocol was also successful for production and selection of multinucleated homokaryons.

## MATERIAL AND METHODS

### Cell culture

The mouse fibroblast cell line NIH/3T3 was purchased from ATCC (CRL-1658); primary bovine fetal fibroblasts (bFFs) were derived from a male fetus of unknown genetic background obtained at an abattoir at gestation day 60. Cells were grown in a 5% CO_2_ in air incubator (Forma series II water jacketed incubator) at 37°C (for NIH/3T3 cells) or 38.5°C (for bFFs) on 10 cm tissue culture dishes (Falcon) in fibroblast medium: Dulbecco’s minimal essential medium (DMEM; Gibco) supplemented with 10% Fetal Bovine Serum (FBS; HyClone) and 50 µg/ml Gentamicin (Lonza). Medium was replaced every two to three days. Subculture was done with TrypLE Express (Gibco) before cells reached 80% confluence. NIH/3T3 cells and bFFs were passage at least twice to ensure proper growth. Primary cell lines used for all experiments were between passages 2 to 6. Mouse embryonic stem (mES) cells from the line C57BL/6 were purchased from ATCC (CRL-1002) and cultured in mES cell medium: DMEM with 15% ES cell-qualified fetal bovine serum (FBS; Gibco), 1X Non-Essential Amino Acids (HyClone), 1X Glutamax (Gibco), 1,500 U/ml of ESGRO (Millipore), 0.55 mM beta-mercaptoethanol (Sigma), 1 μM PD0325901 (PD; Cayman chemical company), 3 μM CHIR99021 (CHIR; Cayman chemical company), and 50 μg/ml Gentamycin (Lonza); mES cells were cultured either on a monolayer of gamma irradiated mouse embryonic fibroblasts (IRR-MEFs; StemGent) or gelatin coated dishes (Sigma). Fused monolayers of NIH/3T3 cells and bFFs were cultured in fibroblast medium, or in serum starvation medium[4]: DMEM, 0.5% FBS, 1% non-essential aminoacids (HyClone), 1% Glutamax (Gibco), and 50 μg/ml Gentamicin. Fused monolayers of co-cultured bFFs and mES cells were incubated in basic ES cell medium: DMEM with 15% ESC-qualified FBS (Gibco), 1X Non-Essential Amino Acids (HyClone), 1X Glutamax (Gibco), 1,500 U/ml of ESGRO (Millipore), 0.55 mM beta-mercaptoethanol (Sigma), and 50 μg/ml Gentamycin (Lonza). Upon selection, hybrids were cultured on IRR-MEFs, ∼200 sorted cells per 24-well plate well, in ES cell medium.

### Homokaryon production

One day before fusion, cells were detached with TrypLE to a monocellular suspension and counted with a hemacytometer. We plated 0.1×10^6^ NIH/3T3 cells or 0.05×10^6^ bFFs in 24-well tissue-culture treated Falcon™ polystyrene flat-bottom microplates (Fisher scientific) and incubated overnight, which generated a ∼95% confluent monolayer 12h later. For fusion, all media, buffers, and reagents were pre-warmed to 37°C, and media changes were performed carefully from the side of the dish to avoid detachment of the cell monolayer. First, we tested the fusogenic effect of (a) 50% PEG 1500 (Roche) (b) 25% PEG 1500 (c) 50% PEG 3000-3700 (Sigma), or (d) 25% PEG 3000-3700. PEG solutions come as a 50% w/v dilution in Hepes; to generate 25% w/v we diluted PEG by adding an equal volume of Dulbecco’s Phosphate-Buffered Saline (DPBS; HyClone). Cells were washed twice with 1 ml DPBS each time before addition of 50 µl of PEG for exactly 2 min at room temperature, followed by two successive washes with DPBS and one wash with DMEM (1 ml each one). All treatments were run in parallel with 1 to 3 wells for each one, and this was replicated four times. For every replicate, one or two wells were treated with all washing steps but no PEG. Cells were incubated for 6h at 37°C, detached using TrypLE, stained with Hoechst 33342, and multinucleated cells counted with a hemacytometer. Next, we fused NIH/3T3 cells using four different PEG treatments, with every treatment replicated using three different volumes of PEG. Treatments were: (a) PEG 1500, (b) PEG 1500 additioned with 10% DMSO, (c) pre-treatment of cells with hypoosmolar buffer for 2 min, followed by PEG 1500, and (d) pre-treatment of cells with hypoosmolar buffer for 2 min, followed by PEG 1500 additioned with 10% DMSO; every treatment was replicated using three different volumes of PEG: 50 µl, 100 µl, and 200 µl. For treatments using hypoosmolar buffer, a wash step using isoosmolar buffer prior to PEG treatment was used. Isoosomolar potassium phosphate buffer (10 mM KH_2_PO_4_, 10 mM K_2_HPO_4_, 1mM MgCl_2_, and 250 mM sucrose, in dH2O) and hypoosmolar potassium phosphate buffer (10 mM KH2PO4, 10 mM K2HPO4, 1mM MgCl2, 75 mM sucrose, in dH2O) were prepared as previously described[22], filter-sterilized, and stored at 4°C before use. Every treatment was replicated four times, with 1 to 3 wells each time. For every replicate, one or two wells were treated with all washing steps but no PEG. Cells were incubated for 12 h at 37°C, detached using TrypLE, stained with Hoechst, and binucleated cells counted with a hemacytometer. Last, we fused bFFs with 200 µl PEG 1500 plus 10% DMSO as described above, and incubated for 12h before analysis.

### Heterokaryon production

Heterokaryons were prepared similarly as described for homokaryons. We seeded 2×10^5^ bFFs in one well of a 24-well plate in basic ES cell medium, and 2 h later 6×10^5^ mESCs were seeded in the same well, without replacing the culture medium. Cells were co-cultured in basic ES cell medium for 4h before fusion. For fusion, monolayers were washed twice with 1 ml DPBS each time before addition of 200 µl PEG 1500 additioned with 10% DMSO (180 µl of 50% PEG 1500 solution plus 20 µl DMSO, mixed and pre-warmed to 37°C) for exactly 2 min at room temperature, followed by two successive washes with DPBS and one wash with basic ES cell medium (1 ml each one). Cells were cultured at 37°C in basic ES cell medium until analysis. A co-culture control (bFF and mESC plated as described but no fusion) was also included in one of the replicates.

### Trypan blue viability assay

Trypan blue (Hyclone) was syringe-filtered and diluted 1:1 in DPBS. Cells were detached using TrypLE and resuspended in 50 µl DPBS containing trypan blue and Hoechst. Suspension was loaded on a hemacytometer and examined under an inverted microscope (Olympus) to determine the percentage of viable cells (clear cytoplasm) versus nonviable cells (blue cytoplasm); Hoechst was used to observe nuclei.

### Indirect immunostaining

Fused monolayers were dissociated to a monocellular suspension with TrypLE (Gibco), centrifuged, resuspended in DNase solution (0.1 mg/ml deoxyribonuclease I in DMEM; Worthington biochemical corporation) and incubated for 15 min at room temperature. Following incubation, solution was filtered through a 100 µM cell strainer (Thermo Fisher). Filtered cells were pelleted by gentle centrifugation and resuspended in DPBS plus 1% heat-inactivated FBS containing antibodies or Hoechst, incubated in the following order: anti-mouse/human CD44 (1:100 dilution, 30 min incubation; Biolegend cat#103001), rabbit anti-rat AlexaFluor® 488 (1:1000 dilution, 20 min incubation; Jackson Immunoresearch Inc., Cat#3125455003), anti-mouse SSEA-1 (1:100 dilution, 30 min incubation; Santa Cruz Biotechnology, sc-21702), and goat anti-mouse Alexa Fluor® 647 (1:2000 dilution, 20 min incubation; Abcam ab150123), and Hoechst (15 min). Between staining steps, cells were pelleted by centrifugation and supernatant removed before resuspending in a new staining solution; incubation was at 4°C in the dark. After the last incubation step, cells were washed and resuspended in DPBS with 1% FBS. Controls (negative (unstained), single colors (CD44/A488, SSEA-1/A647, and Hoechst), secondary antibodies) were always prepared in parallel, following all steps described for samples.

### Flow cytometry

Cells were stained as described and resuspended in DPBS with 1% FBS to a concentration of ∼5×10^6^ cells/ml for ImageStream Mark II (Amnis) and ∼1×10^6^ cells/ml for FACSAria I (BD) analysis. The ImageStream is an imaging flow cytometer that permits visualization (bright field and fluorescence) of cells directly in flow; the ImageStream does not perform sorting. The FACSAria is a sorting cytometer, and parameters identified on the ImageStream can be used to sort cells in the FACSAria. We first analyzed cells using ImageStream to identify location of bicolored, multinucleated heterokaryons. The preprocessing of the images and identification of the area with highest presence of heterokaryons was performed with IDEAS ImageStream Analysis Software. For all replicates, compensation was done with single color stained samples. The obtained parameters were used to sort heterokaryons with the FACSAria.

### Statistical analysis

Data was organized on a Microsoft Excel 2016 spreadsheet and analyzed using JMP (version 13.2.1). We used one- or two-way ANOVA (as noted in the legend below graphs). Multiwell plates run at different time points were accounted as a blocking variable. When statistical significance was confirmed in the ANOVA, differences between groups were determined using t-test or the Tukey-Kramer multiple comparison post-hoc test, when appropriate. P-values <0.05 were considered statistically significant.

## RESULTS

### NIH/3T3 fibroblast monolayers can be fused with 50% PEG 1500 (w/v)

We tested the effect of PEG 1500 and PEG 3000-3700 at two different concentrations (25% and 50%) and counted total and multinucleated cells 6h after fusion treatment (Figure 1A). The majority of the multinucleated cells presented two nuclei. Occasionally, it was possible to observe cells with >2 nuclei but for consistency these were not counted. We found that 50% PEG 1500 and 50% PEG 3000-3700 resulted in 7.28% and 5.58% of binucleated cells, respectively (Figure 1B). We studied the viability of the monolayers after fusion, and observed that NIH/3T3 cells treated with 50% PEG 3000-3700 were not viable after 24h, whereas the remaining treatments appeared viable and were indistinguishable from untreated controls (Figure 1C). Images of untreated cells cultured in parallel are presented for comparison (Figure 1D). Fused monolayers were cultured for four passages; cells treated with 50% PEG 3000-3700 did not recover, whereas the remaining treatments were indistinguishable from the control group. Given that the damage to the cells was not immediately visible, for experiment presented in the next two sections, monolayers were left for 24h in serum starvation medium before fusion percentage and viability were assessed. Serum starvation conditions were necessary to prevent monolayers from overgrowing and detaching from the culture plate.

**Figure 1.**
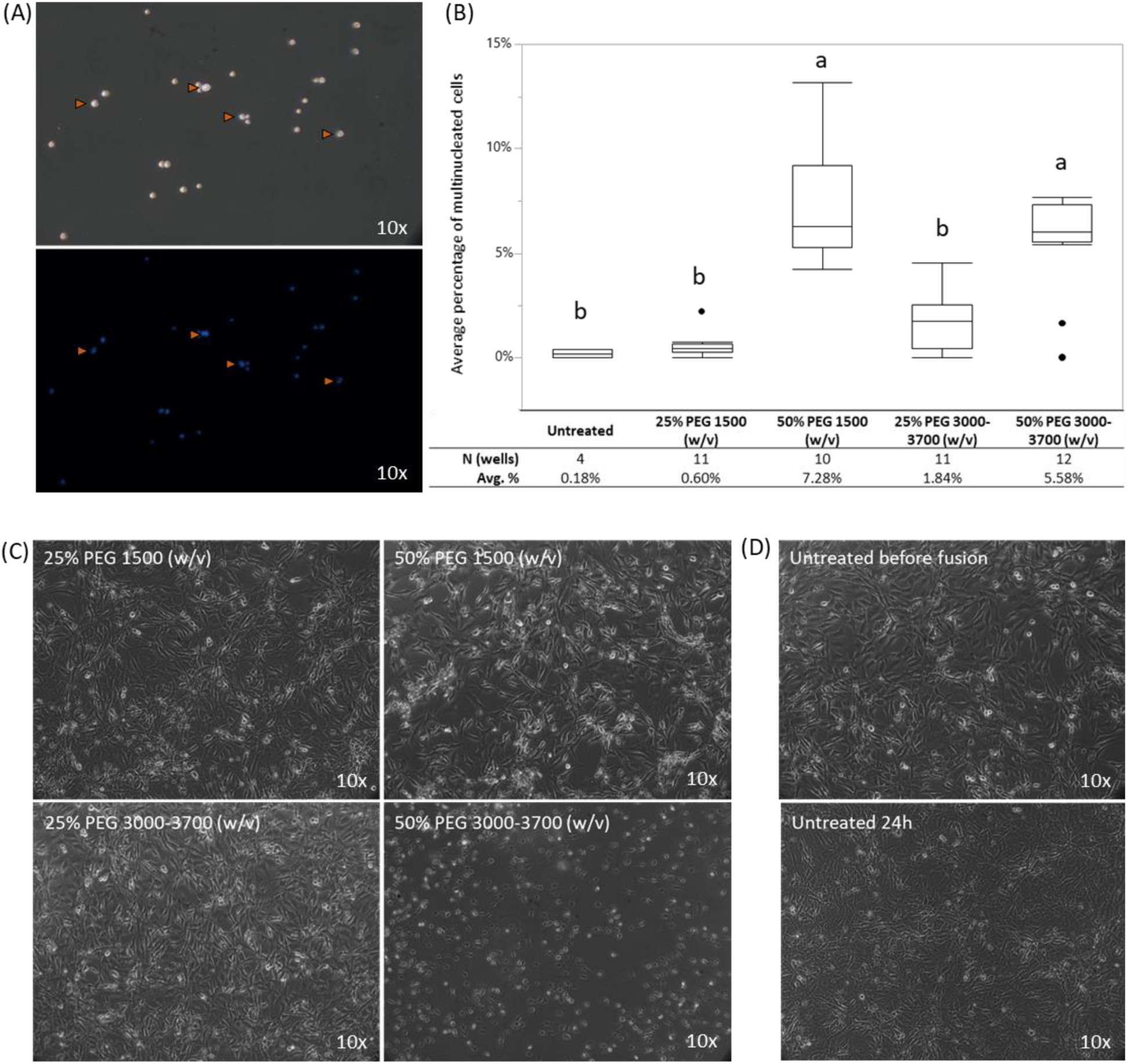
Fusion of NIH/3T3 fibroblasts with four PEG-based fusion treatments. **(A)** Six hours after fusion, cells were detached, stained with Hoechst nuclear stain (blue), and loaded onto a hemacytometer for counting of total and multinucleated cells (orange arrowheads). **(B)** Boxplot representing average percentage of multinucleated cells 6h after treatment; a small number of multinucleated cells also appeared in unfused wells. One-way ANOVA followed by Tukey’s HSD test. Bars are SEM. **(C)** After fusion treatment, cells were left in culture for 24h. Cells exposed to 50% PEG 3000-3700 did not appear viable 24h after treatment. **(D)** Near confluent monolayer of NIH/3T3 cells before fusion (top) and equivalent monolayer after 24h (bottom).

### Addition of 10% of DMSO increases fusion efficiency in NIH/3T3 cells

Next, we determined if an increase of PEG volume, addition of 10% DMSO to the PEG mixture, and/or pre-treatment of monolayers with hypoosmolar buffer, had an impact on cell fusion. PEG is a viscous solution and small volumes can be difficult to pipet, with the risk of administering a suboptimal volume of PEG to the cells; because of this, we increased the volume of PEG used in the same culture well. We observed that, regardless of treatment, PEG volume had no meaningful statistical significance over the resulting fusion percentage (Table 1). Incorporation of 10% DMSO to the PEG mixture has been described to increase the frequency of cell fusion in some cell types[23].We observed that addition of 10% DMSO had a positive effect, resulting in 11.71% yield of multinucleated cells (Figure 2A). Hypoosmolar buffer is used to increase cell volume and therefore increment the surface exposed for fusion. However, pre-treatment of monolayers with hypoosmolar buffer affected cell adhesion (Figure 2B), causing detachment of part of the monolayer in 26% and 37% of the wells exposed to this treatment (Table 2). Due to its higher efficiency and easiness to pipette, subsequent experiments were performed using 200 µl of 50% PEG 1500 with 10% DMSO, for every individual well of a 24 well plate.

**Table 1.** Average percentage of multinucleated cells after treatment with 50% PEG 1500 and three variations of this method, each one tested with three different volumes. Data was analyzed with two-way ANOVA for treatment and volume of PEG. Neither volume (P=0.84) or the interaction between treatment and volume (P=0.26) were statistically significant. Treatment was significant and is shown in Figure 2A.

| Pre-treatment | PEG | Additive | PEG solution<br>final volume | N (wells) | Mean<br>fusion % |
| --- | --- | --- | --- | --- | --- |
|  | 50% PEG 1500 (w/v) |  | 50 µl | 8 | 4.12% |
|  |  |  | 100 µl | 9 | 4.14% |
|  |  |  | 200 µl | 8 | 4.29% |
|  | 50% PEG 1500 (w/v) | 10% DMSO | 50 µl | 8 | 10.34% |
|  |  |  | 100 µl | 9 | 13.71% |
|  |  |  | 200 µl | 8 | 11.08% |
| Hypoosmolar buffer | 50% PEG 1500 (w/v) |  | 50 µl | 5 | 4.96% |
|  |  |  | 100 µl | 5 | 4.31% |
|  |  |  | 200 µl | 7 | 5.45% |
| Hypoosmolar buffer | 50% PEG 1500 (w/v) | 10% DMSO | 50 µl | 7 | 9.04% |
|  |  |  | 100 µl | 7 | 7.30% |
|  |  |  | 200 µl | 6 | 6.75% |
| Untreated |  |  |  | 7 | 0.29% |
| Isoosmolar buffer wash step |  |  |  | 7 | 0.37% |

**Table 2.**
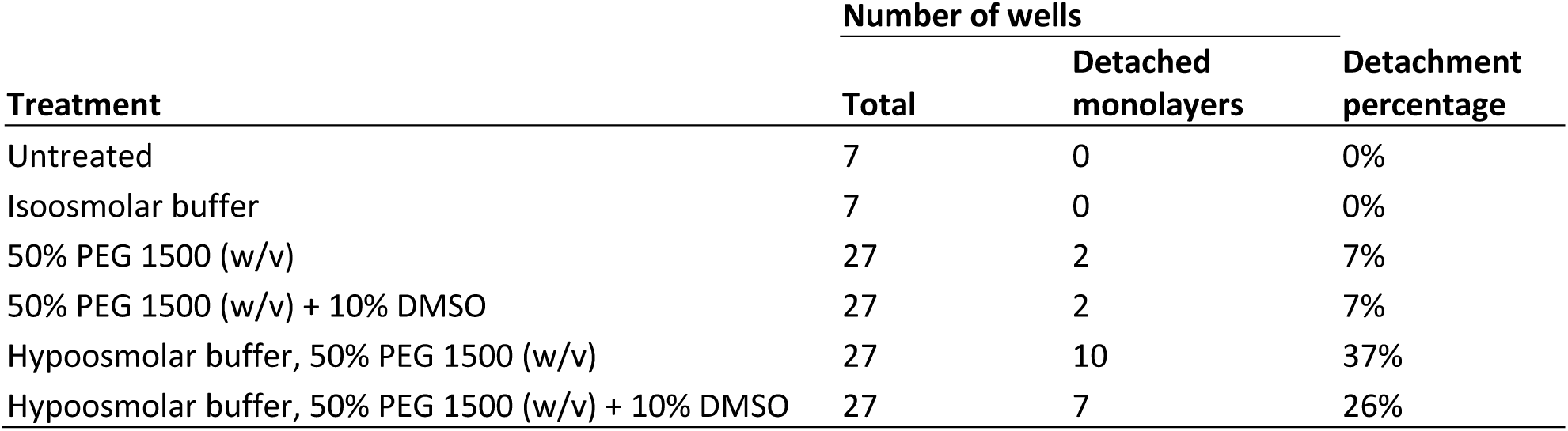
Detachment of NIH/3T3 cell monolayers after different treatments. A monolayer was considered to be detached when more than ∼40% of the cells were lost (by visual inspection).

| Treatment | Number of wells |  | Detachment percentage |
| --- | --- | --- | --- |
|  | Total | Detached monolayers |  |
| Untreated | 7 | 0 | 0% |
| Isoosmolar buffer | 7 | 0 | 0% |
| 50% PEG 1500 (w/v) | 27 | 2 | 7% |
| 50% PEG 1500 (w/v) + 10% DMSO | 27 | 2 | 7% |
| Hypoosmolar buffer, 50% PEG 1500 (w/v) | 27 | 10 | 37% |
| Hypoosmolar buffer, 50% PEG 1500 (w/v) + 10% DMSO | 27 | 7 | 26% |

**Figure 2.**
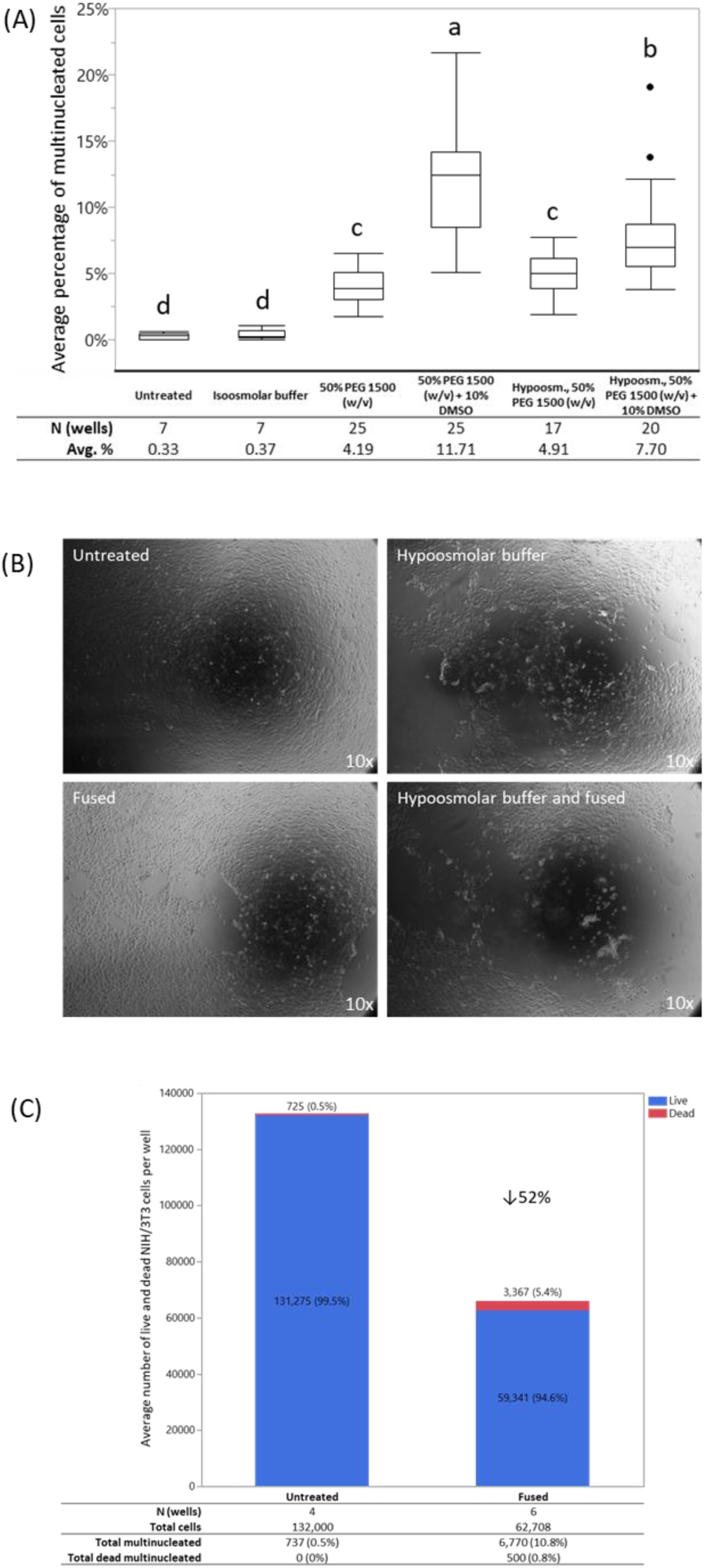
Fusion of NIH/3T3 cells with 50% PEG 1500 and three variations of this method. **(A)** Average percentage of multinucleated cells after different fusion treatments. Controls included untreated and isoosmolar buffer treatment. Statistical significance was determined by two-way ANOVA for treatment and volume of PEG. Only treatment was statistically significant (P<0.05); significant groups were determined by Tukey HSD test. Bars indicate SEM. **(B)** Representative images of NIH/3T3 monolayers 6h after treatment with hypoosmolar buffer only (top right), 50% PEG 1500 (bottom left), and hypoosmolar buffer followed by 50% PEG 1500 (bottom right); untreated control (top left) also included. **(C)** Trypan blue viability assay of NIH/3T3 cells 24h after fusion treatment. Stacked bars indicate average number of viable and dead NIH/3T3 cells per well. Cell number reduced by 52% in fused wells compared to untreated wells. Percentage of dead multinucleated cells (0.8%) was lower than the general percentage of dead cells (5.4%).

Cells exposed to 50% PEG 1500 plus 10% DMSO appeared viable 24 h after treatment. Monolayers were harvested and viability determined using Trypan blue. Cell mortality was higher in the fused group compared to PEG-free control (5.4% and 0.5%, respectively). Nevertheless, mortality was not increased in multinucleated versus single nucleated cells (Figure 2C). Both fused and control cells proliferated similarly for three passages, at which point the cultures were ended.

### Fusion of bFFs with 50% PEG 1500 plus 10% DMSO produces a similar percentage of multinucleated cells as in mouse NIH/3T3 fibroblasts

We fused near confluent bFF monolayers with 200 µl 50% PEG 1500 plus 10% DMSO, and counted total and multinucleated cells 24h after treatment (Figure 3A). Fusion treatment resulted in 11.05% of multinucleated cells, in contrast to the 2.81% found in the control group (Figure 3B). Cell mortality was higher in the fused group compared to PEG-free control (4.9% and 0.7%, respectively). Nevertheless, mortality was not increased in multinucleated versus single nucleated cells (Figure 3C). Both fused and control cells proliferated similarly for three passages, at which point the cultures were ended.

**Figure 3.**
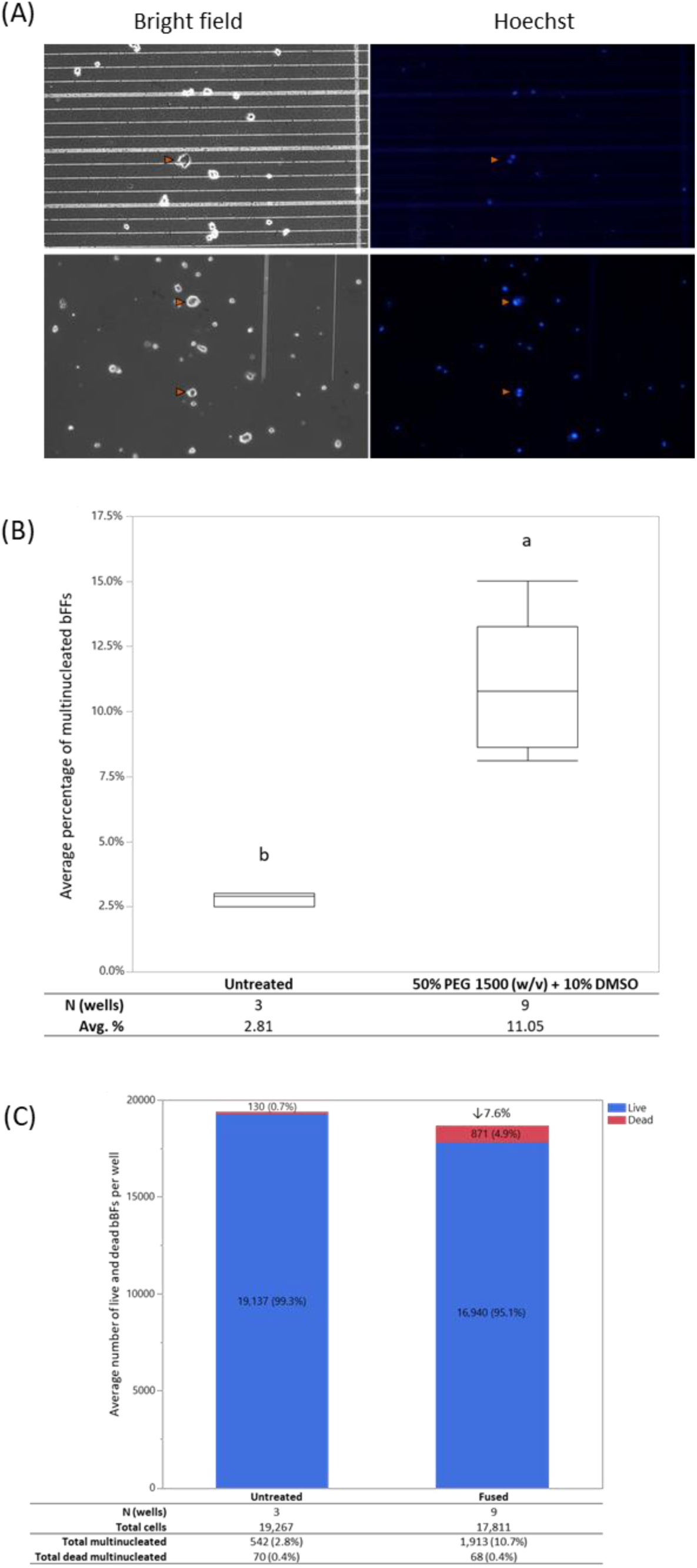
Characterization of bFFs after fusion with 50% PEG 1500 plus 10% DMSO. **(A)** Following fusion treatment, cells were left overnight in serum starvation medium; cells were then detached, stained with Hoechst, and loaded onto a hemacytometer to count for total and multinucleated cells (orange arrowheads). **(B)** Average percentage of multinucleated bFFs after fusion treatment. Statistical significance was determined by one-way ANOVA followed by t-test. **(C)** Trypan blue viability assay of bFFs 24h after fusion. Stacked bars indicate average number of viable and dead bFFs per well. Cell number reduced by 7.6% in fused wells compared to untreated wells. Percentage of dead multinucleated cells (0.4%) was lower than the general percentage of dead cells (4.9%).

### Interspecies heterokaryons can be identified and selected using ImageStream and FACSAria

Fused monolayers of co-cultured bFFs and mESCs were stained and immediately analyzed using ImageStream, to identify the area in the histogram that contains potential heterokaryons (Figure 4A-D). Figure 4E shows a representative screenshot of the area in the histogram where the majority of the heterokaryons were found. Heterokaryons represented >50% of the cell population in this gate, but false positive cell aggregates were also observed. Distinct nuclei were observed in heterokaryons sampled at 24 and 48h (Figure 4F and G), whereas the double positive heterokaryons at 72h presented enlarged and/or fragmented nuclei (Figure 4H). The parameters identified on the ImageStream were extrapolated to the FACSAria flow cytometer (Figure 5). The percentage of double positive cells in the total sample was ∼0.1% for almost all time points, and was constant in both replicates (Table 3).

**Table 3.** Double positive cells observed with FACSAria I, expressed as percentage of total cells.

|  | <u>% of total double positive cells</u> |  |  |
| --- | --- | --- | --- |
|  | 24h | 48h | 72h |
| Rep1 | 0.3 | 0.1 | 0.1 |
| Rep2 | 0.1 | 0.1 | 0.1 |

**Figure 4.**
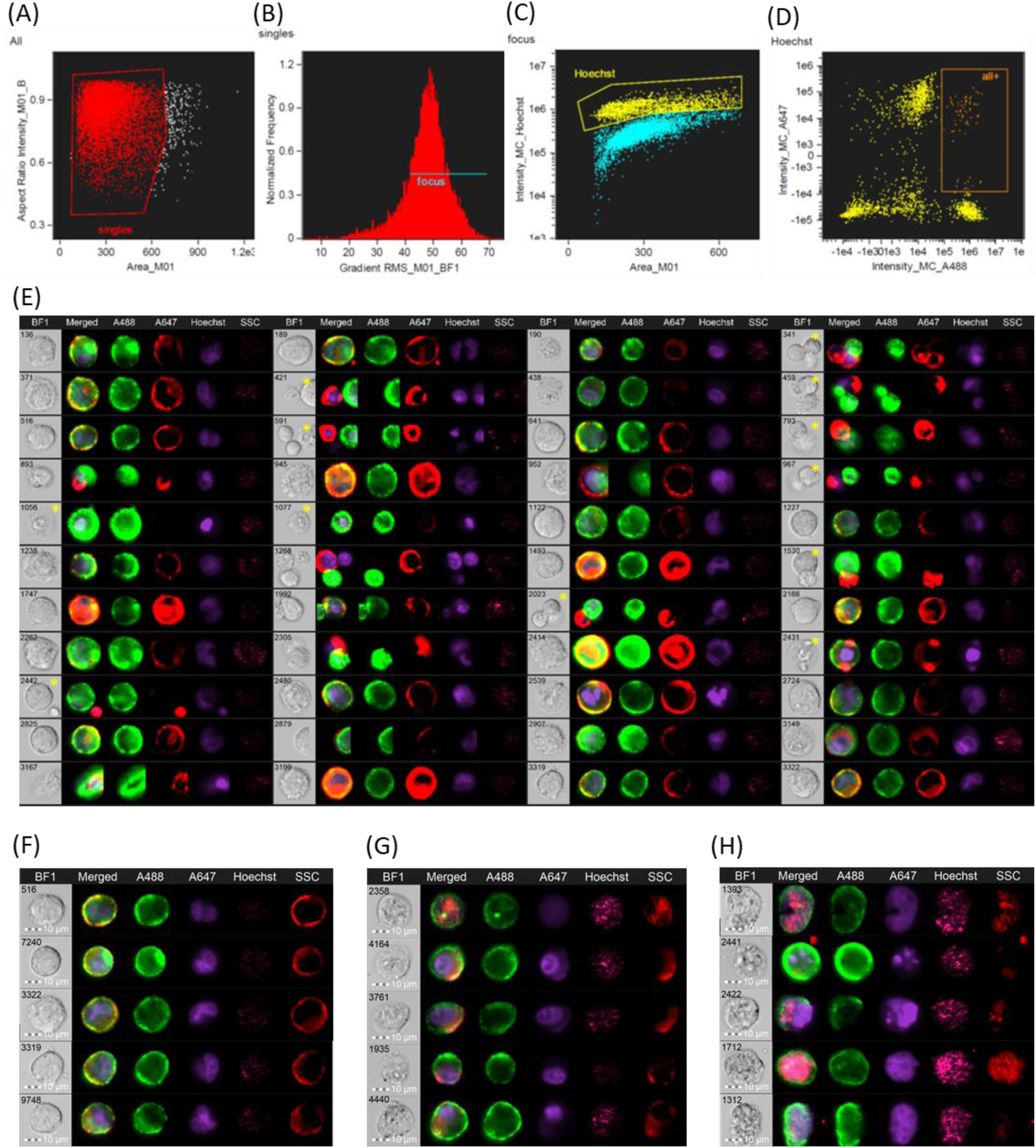
Identification of heterokaryons using the ImageStream Mark II imaging flow cytometer. Only single cells **(A)** in focus **(B)** were considered. The remaining cells were plotted as intensity of Hoechst vs. cell area in the bright field image **(C)**, and a population of high intensity of Hoechst was further selected and plotted as intensity of Alexa 488 vs. Alexa 647 **(D)**. In this scatterplot, a population of high A488 and high A647 was found to contain a majority of Heterokaryons (orange rectangle); a representative image of the cells found in this population is shown in **(E)**. It was still possible to find false positive cells in this subpopulation (yellow asterisks in bright field image) but it was not possible to gate these events out. The parameters identified on the ImageStream were used to sort heterokaryons in the FACSAria. Representative images of heterokaryons found **(F)** 24h, **(G)** 48h, and **(H)** 72h after fusion are shown.

**Figure 5.**
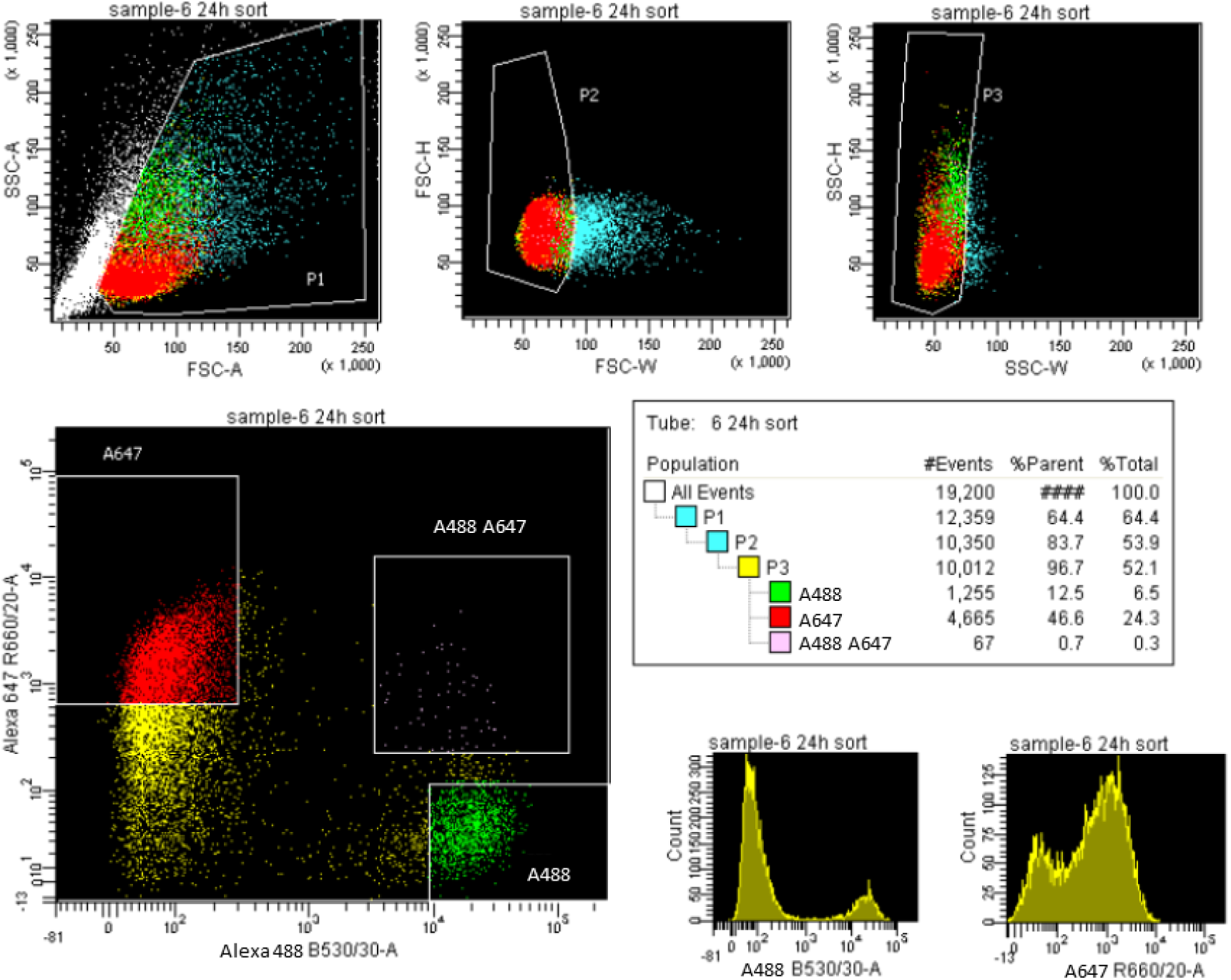
Representative image of gating strategy used to select heterokaryons on FACSAria I. Heterokaryons are present in the top right quadrant of the A647 vs A488 scatterplot.

### Homokaryons can be identified and selected using ImageStream and FACSAria

Homokaryons were also were identified based on cell size and intensity of cell type-specific dye. Figure 6 shows the workflow used to identify bFF homokaryons by ImageStream, which were then selected using FACSAria (Figure 7).

**Figure 6.**
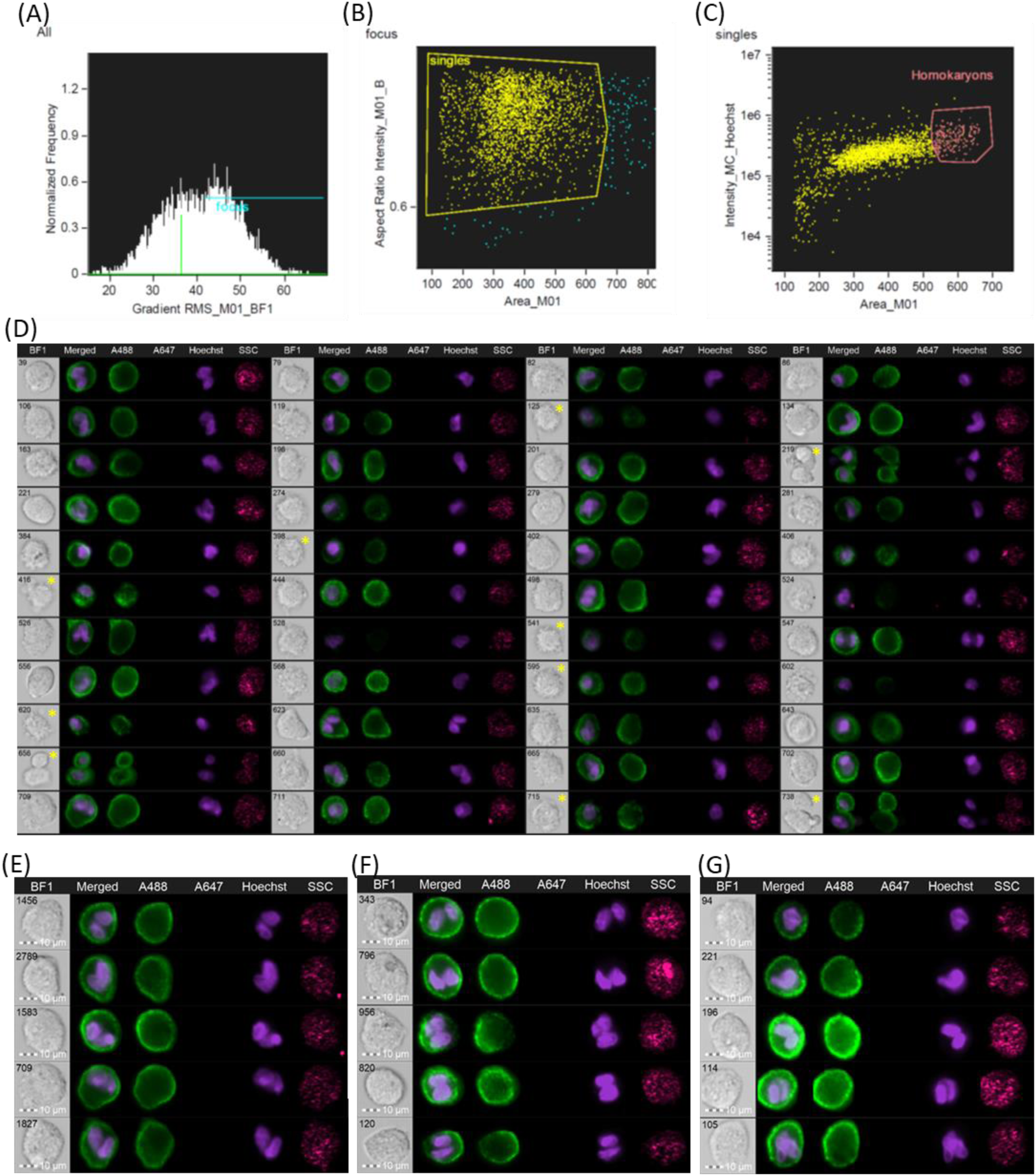
Identification of homokaryons using ImageStream Mark II imaging flow cytometer. Similar to heterokaryons, only single cells in focus were considered **(A and B)**, and the remaining cells were plotted as intensity of Hoechst versus cell area **(C)**; in that plot, a section containing a majority of homokaryons was found, as shown in **(D)**. It was still possible to find false positive cells in the subpopulation containing a majority of homokaryons (yellow asterisk in bright field image) but it was not possible to gate these events out any further. The parameters identified on the ImageStream were used to sort homokaryons in the FACSAria. Representative images of homokaryons found **(E)** 24h, **(F)** 48h, and **(G)** 72h after fusion are shown.

**Figure 7.**
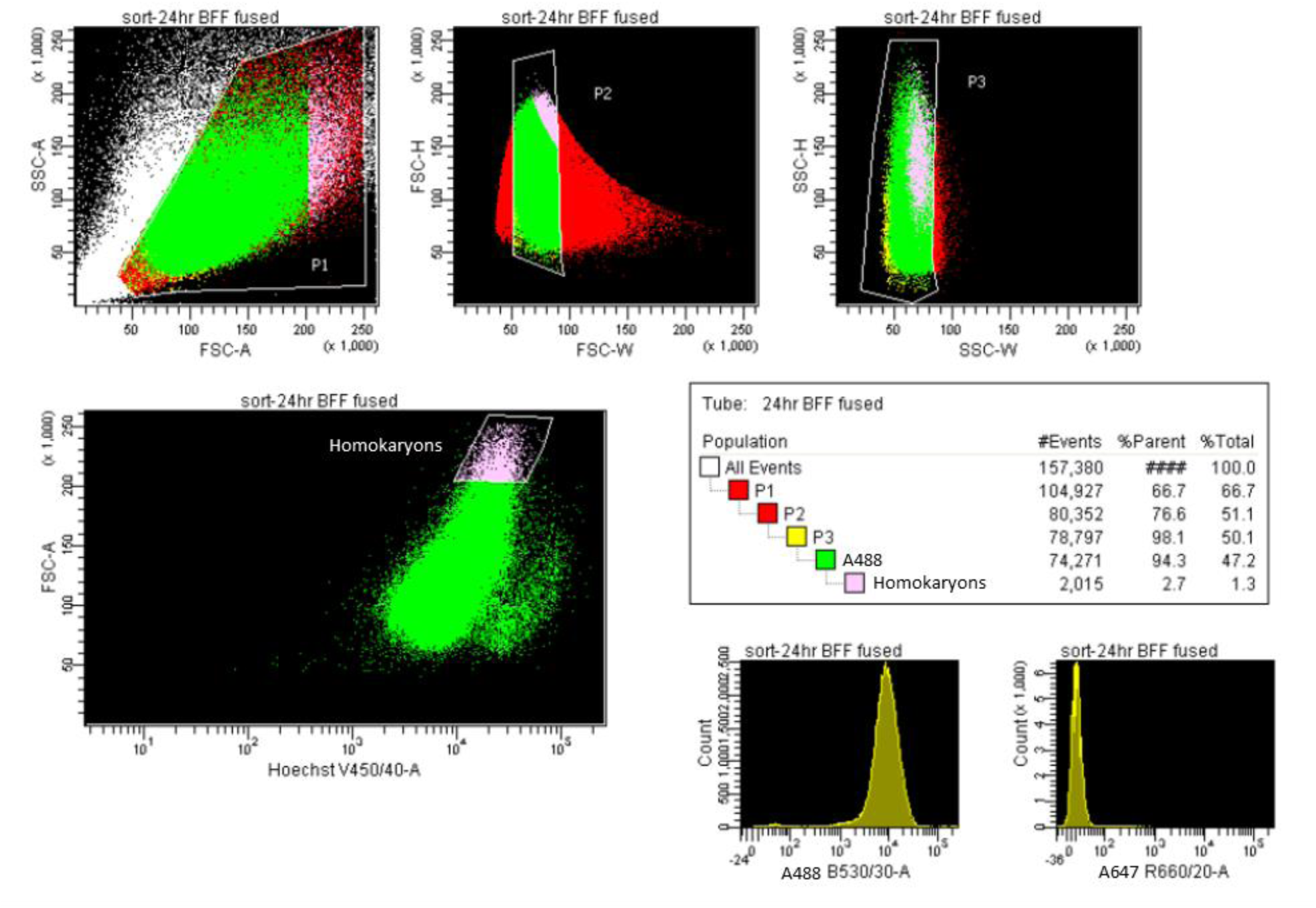
Representative image of gating strategy used to select bFF homokaryons using FACSAria I.

### Hybrids can be produced from sorted heterokaryons

We selected 600 double-positive heterokaryons and 600 bFF homokaryons at 24, 48, and 72h after fusion using FACS, and cultured on IRR-MEFs. Colonies were visible in heterokaryon cultures as early as 24h and were counted six days after plating, with a few isolated colonies also observed in homokaryon cultures (Table 4). We observed two distinct types of colony morphologies at all time points: tightly packed colonies with a defined border, and colonies with an irregular border surrounded by a translucent, flat looking “halo” of cells (Figure 8). Interestingly, hybrid colonies obtained at different time points after fusion presented different proportions these two of morphologies, with number of colonies with irregular borders increasing over 48 and 72h.

**Table 4.** Number of colonies observed after six-day culture of heterokaryons and homokaryons selected at different time points after fusion. Two distinct colony morphologies were observed at each time point. Percentage indicated is based on the 600 cells initially sorted

| Morphology | 24h |  |  |  | 48h |  |  |  | 72h |  |  |  |
| --- | --- | --- | --- | --- | --- | --- | --- | --- | --- | --- | --- | --- |
|  | Heterok. |  | Homok. |  | Heterok. |  | Homok. |  | Heterok. |  | Homok. |  |
|  | # | % | # | % | # | % | # | % | # | % | # | % |
| Irregular border | 14 | 2.3% | 1 | 0.2% | 33 | 5.5% | 0 | 0.0% | 63 | 10.5% | 0 | 0.0% |
| Defined border | 14 | 2.3% | 0 | 0.0% | 26 | 4.3% | 0 | 0.0% | 45 | 7.5% | 2 | 0.3% |
| Total colonies | 28 | 4.7% | 1 | 0.2% | 59 | 9.8% | 0 | 0.0% | 108 | 18.0% | 2 | 0.3% |

**Figure 8.**
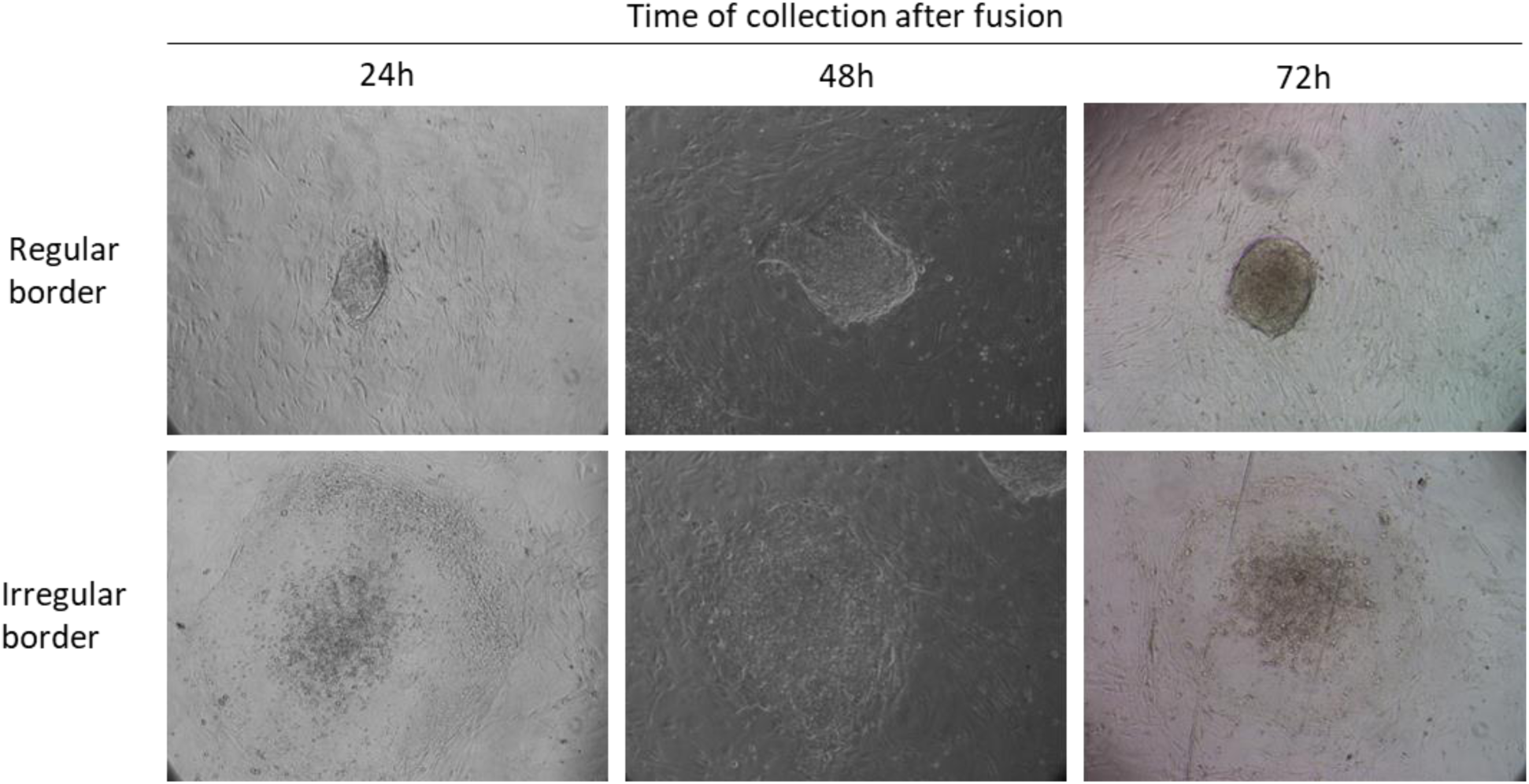
Representative images of colonies observed after six-day culture of heterokaryons selected at different time points after fusion. Two characteristic morphologies were observed in all groups. Images are 10X.

## DISCUSSION

The interspecies cell fusion approach holds great potential for the study of nuclear reprogramming. Here, we sought to establish a method to consistently produce a high percentage of bi-species multinucleated cells. Initially, we tested several conditions previously described in the literature and tested on cell from the same cell type (NIH/3T3 or bFFs). Although cell fusion has been used for decades, data on the percentage of fused cells obtained after treatment is not always available. Due to technological limitations in the past, early cell fusion work studied hybrid colonies, which form days after the initial fusion event[24]. It is known that not all fusion products form hybrids, making it difficult to estimate how many cells initially fused. Efficiency of cell fusion also varies depending on the cell type(s), the fusogen, and the method used to determine fusion. For example, Blau *et al.* (1983) fused human amniocytes and mouse myotubes with PEG, and observed an average of 73% of heterokaryon formation by visual inspection of multinucleated cells[15]. In contrast, Brady *et al.* (2013) used PEG to fuse GFP^+^ mouse ESCs and dsRed^+^ human primary fibroblasts, and used FACS to select for double positive heterokaryons, obtaining 1.16% of heterokaryons on a first sort, which was later increased to 51.6% after a second sort and 77.8% after enrichment[19]. Due to the heterogeneity of the available data, as well as the different methods used to determine fusion efficiency in early experiments, we used direct microscopic visualization of multinucleated cells to avoid the possibility of instrumental error.

Using PEG alone, the highest percentage of multinucleated NIH/3T3 cells was obtained using either 50 µl of 50% PEG 1500 or 50% PEG 3000-3700 per well in a 24-well plate for 2 min. Work in amphibians by Broyles et al. (2006) found that PEG-mediated fusion is a gradual process, with peak number of heterokaryons found after 4 – 6 hours[25]. Initially, we counted the number of multinucleated cells 6h after fusion treatment. However, cells treated with PEG 3000-3700 were not viable 24h after treatment, indicating that cell mortality might not be visible in the short term. Therefore, for the remainder of the study we incubated the cells for 24h before counting multinucleated cells. To prevent nuclear fusion during incubation, in our initial homokaryon study we cultured the cells in low serum conditions, which prevents the majority of the cells from resuming cell cycle. PEG has also been associated with toxicity by others; we therefore recommend assessing the sensitivity of the cell types being used.

Next, we tested if the increase of PEG volume, the addition of 10% DMSO, and/or the pre-treatment of the cells with hypoosmolar buffer, could increase cell fusion in NIH/3T3 cells. Depending on the cell types being fused, the addition of 10% DMSO to the PEG mixture increases the frequency of cell fusion, as has been described for mouse spleen cells and mouse ESCs[26]. However, the addition of DMSO appeared to have no effect over frequency of fusion nor viability when fusing mouse C2C12 muscle cells with human amniocytes[15]. In our study, we found that addition of 10% DMSO to the PEG solution had a positive effect over the number of multinucleated NIH/3T3 cells, resulting in a 62% increase of fusion. Volume ranging from 50 µl to 200 µl had no significant impact over the percentage of fused cells. We also tested the effect of pre-treating the cells with hypoosmolar buffer, a procedure commonly used to increase cell volume and therefore incrementing the surface exposed for fusion, when inducing fusion in cell suspensions by electrofusion. However, we found that pre-treatment with hypoosmolar buffer had no beneficial effect when fusing cells growing on a culture plate. Moreover, it caused detachment of the cellular monolayer, indicating that this treatment may not be adequate to fuse cells growing on monolayers.

It has been described that different cell types can have different fusion potentials when exposed to PEG[14], Therefore, toxicity, molecular weight, and concentration of the PEG solution, as well as the length of the treatment, should be assessed for every cell type. Nevertheless, when we fused mouse NIH/3T3 fibroblasts or bFFs with 200 µl of 50% PEG 1500 plus 10% DMSO, we obtained similar percentages of multinucleated cells. Mortality was also similar for both cell types. For all treatments in our study, we used the same volume and dish size. Although the treatment can in theory be scaled, plate size-dependent events such as meniscus effect[27] or uneven cell densities caused by swirling of the media due to manipulation or vibrations inside the incubator or biosafety cabinet, can potentially have an effect over the effectivity of the treatment. Therefore, we recommend characterizing the efficiency when adapting this protocol to a different culture vessel.

For downstream applications, multinucleated cells have to be isolated from the bulk of the unfused cells; in addition, heterokaryons have to be distinguished from multinucleated homokaryons. Before developing the method described here, we attempted selection of heterokaryons using FACS alone, manual selection with a micromanipulator, or the use of lipophilic tracers, fluorescent nanoparticles and constitutive lentiviral vectors[28]; these methods were either unspecific, required an extended operation time, and/or negatively affected viability of the cells. Therefore, to specifically select an enriched population of heterokaryons while preserving viability, we used a combination of imaging flow cytometry and FACS to select interspecies heterokaryons, stained with cell type specific fluorescent antibodies. Cells were first analyzed using ImageStream, to identify a subpopulation of cells that had presence of both fluorophores and two nuclei. The parameters were then used to sort this subpopulation of cells. We found that ∼0.1% of the cells were double positive on a first sort. To better preserve viability of the cells, we did not attempt enrichment sorts. Sorted cells can be placed in culture or used immediately for downstream applications such as nucleic acid isolation.

## CONCLUSIONS

We present a method to consistently produce bi-species somatic/pluripotent (bFF/mES cell) heterokaryons, using 50% PEG 1500 plus 10% DMSO. Multinucleated cells were detected by indirect immunofluorescence in live cells, using cell-type specific markers. We used ImageStream imaging flow cytometry to identify the cell population of interest, and then select these cells using FACS. With this method, we were able to obtain an enriched population of double positive cells, which accounted for ∼0.1% of the total cell population. The selected heterokaryons have been used for nucleic acid extraction and are also capable of producing hybrid colonies. The method described here could be adapted to other cell types and species, and aid in the understanding of nuclear reprogramming mechanisms. Our method is also suitable for production of as same species homokaryons.

